# Cuban history of CRF19 recombinant subtype of HIV-1

**DOI:** 10.1101/2021.02.15.431210

**Authors:** Anna Zhukova, Jakub Voznica, Miraine Dávila Felipe, Thu-Hien To, Lissette Pérez, Yenisleidys Martínez, Yanet Pintos, Melissa Méndez, Olivier Gascuel, Vivian Kouri

**Author notes:** co-corresponding authors (AZ), (OG), (VK). OG: current address. MDF: current address.

## Abstract

CRF19 is a recombinant form of HIV-1 subtypes D, A1 and G, which was first sampled in Cuba in 1999, but was already present there in 1980*s*. CRF19 was reported almost uniquely in Cuba, where it accounts for ~25% of new HIV-positive patients and causes rapid progression to AIDS (~ 3 years).

We analyzed a large data set comprising ~ 350 *pol* and *env* sequences sampled in Cuba over the last 15 years and ~ 350 from Los Alamos database. This data set contained both CRF19 (~ 315), and A1, D and G sequences. We performed and combined analyses for the three A1, G and D regions, using fast maximum likelihood approaches, including: (1) phylogeny reconstruction, (2) spatio-temporal analysis of the virus spread, and ancestral character reconstruction for (3) transmission mode and (4) drug resistance mutations (DRMs). This allowed us to acquire new insights on the CRF19 origin and transmission patterns. We showed that CRF19 recombined between 1966 and 1977, most likely in Cuban community stationed in Congo region. We further investigated CRF19 spread on the Cuban province level, and discovered that the epidemic started in 1970*s*, most probably in Villa Clara, that it was at first carried by heterosexual transmissions, and then quickly spread in the 1980*s* within the “men having sex with men” (MSM) community, with multiple transmissions back to heterosexuals. The analysis of the transmission patterns of common DRMs showed mostly acquired drug resistance rather than transmitted one.

Our results show a very early introduction of CRF19 in Cuba, which could explain its local epidemiological success. Ignited by a major founder event, the epidemic then followed a similar pattern as other subtypes and CRFs in Cuba. The reason for the short time to AIDS remains to be understood and requires specific surveillance, in Cuba and elsewhere.

**Author summary:** CRF19 is a recombinant form of HIV-1, which causes rapid progression to AIDS (~ 3 years versus 5 – 10 years for other subtypes and CRFs). CRF19 is reported almost uniquely in Cuba, where it is highly prevalent (~ 25%) among newly detected HIV-1 patients. In this study, we found that CRF19 most likely recombined around the 1970*s* in the Cuban community that was stationed in Democratic Republic of the Congo and Angola at that time. It was introduced very early into the Cuban province of Villa Clara, from where it had several introductions to La Habana in the 1980*s* and then further spread to other Cuban provinces. The CRF19 epidemic most probably started with heterosexual transmissions, followed in the 1980*s* by multiple introductions into “men having sex with men” (MSM) community, followed by multiple transmissions back to heterosexuals (often females). The early introduction of CRF19 into Cuba most likely explains its success, not observed in other parts of the world. However, importantly, its rapid progression to AIDS makes it crucial to survey CRF19 sub-epidemics not only in Cuba, but also in other parts of the world having regular exchanges with Cuba.

## Introduction

For HIV-1, more than 100 recombining subtypes have been detected [1]. They have varying epidemiological features, such as pathogenesis and transmission patterns [2]. CRF19 is a recombinant form of HIV-1 subtypes D, A1 and G. It was first detected in Cuba in 1999 when four epidemiologically unrelated individuals (diagnosed in 1997) were sampled [3]. Casado *et al.* [3] proposed an African origin for CRF19 explained by its homology to an AG intersubtype recombinant from Cameroon and a subtype D virus from Gabon. However, while CRF19 has maintained a very low prevalence in the African continent through all these years, the virus has spread successfully in Cuba, placing it among the most prevalent circulating subtypes (24% of all sampled sequences by 2017). While CRF19 identification and first sequences were obtained from patients diagnosed in 1997 [3], a retrospective analysis performed by Perez *et al.* [4] exposed that this recombinant was already infecting Cuban patients who were diagnosed between 1986 – 1990, which evidences an early introduction in the outbreak of the epidemic. Potential central African ancestry of CRF19 could be explained by presence of numerous Cuban military and civilian personnel in several sub-Saharan African countries (the Democratic Republic of the Congo (DRC), Angola, Ethiopia) from 1960s to 1980s [5]. However, CRF19 was never reported in sub-Saharan Africa.

In Cuba, CRF19 was found to cause particularly rapid progression to AIDS (*<* 3 years after seroconversion, versus 5 to 10 years for other subtypes and CRFs [6]) and is associated with drug-resistance [7] in many newly infected patients. Several subsequent introductions related to Cuban samples were then reported in Spain (with one limited outbreak in southern Spain, with mild pathogenicity) [8–10] and Tunisia [11]. Understanding the origin, transmission and drug resistance acquisition of such aggressive subtype is therefore key to impede its further spread in other regions of the world.

Delatorre and Bello [12] performed a maximum likelihood (ML) phylogeographic study of CRF19 and D sequences (as CRF19 is subtype D in the *pol* fragment analysed) from Los Alamos HIV Sequence database. Their data set contained 160 Cuban CRF19 sequences sampled between 1999 and 2011, 3 European CRF19 sequences and 2 000 subtype D sequences from Africa (1996-2011). They discovered that all Cuban CRF19 sequences branched in a highly supported monophyletic sub-cluster that was nested within subtype D *pol* sequences of central African origin. This indicates a common recombined ancestor from which CRF19 spread further. The 3 CRF19 isolates in Europe branched within the Cuban clade, indicating a transmission from Cuba to Europe. They also performed a Bayesian analysis to estimate the time of the most recent common ancestor (MRCA) of the Cuban clade: The median of the estimated date was 1987 (confidence interval (CI) 1983 – 1991).

The goal of this study is to better understand the geographic origin and transmission patterns of CRF19, especially regarding transmission mode and drug resistance, using the new data from Cuba and international databases. New CRF19 and D-infected patients were detected and sequenced in Cuba and worldwide since the study of Delatorre and Bello [12]. Moreover, additional information can be extracted by performing similar studies based on the parts of CRF19 that correspond to subtypes A1 and G. Importantly, with more fine-grained geographic data available for some of the Cuban sequences (e.g. provinces of diagnostics) the CRF19 spread within the country can now be elucidated. Similarly, as for many newly sampled Cuban sequences the information on the risk group is available, it can be used to reconstruct the transmission mode scenarios to learn how CRF19 spread between and within individual communities. Lastly, as transmitted drug resistance (TDR) is becoming predominant in several countries (e.g. UK [13]) and HIV transmission might be further impacted by drug resistance, we also investigated drug resistance mutation spread of CRF19.

We performed maximum likelihood phylogenetic analyses using fast and easy-to-use software, to investigate the spread of the virus, its mode of transmission and drug resistance patterns. Our new Cuban data set contains rich metadata (diagnostics dates, sampling dates, risk groups, province of diagnostics, etc.), from which we obtained spatio-temporal predictions for worldwide and local CRF19 spread as well as its mode of transmission among risk groups in Cuba. The article is organized as follows: we first describe the data set and the insights on CRF19 origin and transmission patterns that we acquired in this study. We then detail the phylogenetic methods used for the temporal, phylogeographic, transmission mode and drug resistance analyses, which could be used for other HIV-1 CRFs or subtypes, and other viruses.

## Results

### Data sets

We combined the data from two sources: a Cuban data set (from now on referred as CU) and the 2018 HIV-1 compedium genome alignment from the Los Alamos HIV database (LA) [14], sampled between 1983 and 2018. LA contained 3 884 sequences of all M-group HIV subtypes and recombinants that were curated. We chose this smaller but curated data set in order to avoid noisy and erroneous data (partial sequences, erroneous date, subtype or country annotations for non-curated sequences, etc.) that could perturb our analyses. CU contained 1 674 partial *pol* and 164 partial *env* sequences (of various subtypes) recently sampled in Cuba, which we subtyped and aligned against LA as described in Materials and methods. We created three data sets out of the combined LA+CU alignment for CRF19 analyses (described in more detail in Table 1):

1. *D+CRF19* – the D-part of CRF19 (C-part of *gag*, PR, RT and *nef*, breakpoints from [3]) for 386 sequences of subtypes D and CRF19 (276 sequences), from CU and LA.
2. *A1+CRF19* – the A1-part of CRF19 (N-part of *gag*, Integrase, *env*) for 218 sequences of subtypes A1 and CRF19 (31), from CU and LA.
3. *G+CRF19* – the G-part of CRF19 (*vif*, *vpr*, *vpu* and C-part of *env*) for 84 sequences of subtypes G and CRF19 (5), from LA (as these parts of the HIV genome were not present in CU).

**Table 1.** Data set description. Number of sequences from different locations, subtypes and sources. LA data contained 6 sequences from Cuba: 3 of subtype G and 3 CRF19.

| data set | sampling dates | total num. | total |  | CRF19 |  |
| --- | --- | --- | --- | --- | --- | --- |
|  |  |  | CU | LA | CU | LA |
| *D+CRF19* | 1983 – 2018 | 386 | 310 | 76 | 271 | 5 |
| *A1+CRF19* | 1985 – 2018 | 218 | 42 | 176 | 26 | 5 |
| *G+CRF19* | 1987 – 2016 | 84 | 0 | 84 | 0 | 5 |

| data set | Africa | Americas | Asia | Europe | Oceania |
| --- | --- | --- | --- | --- | --- |
| *D+CRF19* | UG(36), CD(8), CM(4), ZA(4), KE(3), TD(2), TZ(2), SN(1) | CU(313), BR(2), US(1) | YE(2), CY(1) | GB(4), ES(2), SE(1) |  |
| *A1+CRF19* | KE(63), TZ(24), UG(18), RW(9), CM(3), ZA(2), CD(1) | CU(45) | PK(15), CY(13), IN(5) | SE(10), ES(5), GB(4) | AU(1) |
| *G+CRF19* | NG(28), CM(16), CD(4), KE(3), GH(1), GW(1), ZA(1) | CU(6) | CN(3) | ES(9), PT(4), GB(3), RU(2), BE(1), SE(1) |  |

### Phylogenetic trees

For each data set, we reconstructed phylogenetic trees as described in Materials and methods and found that all the CRF19 sequences formed a unique cluster (see colourstrips in Fig 1). For *A1+CRF19* and *D+CRF19* data sets the CRF19 cluster was placed within the A1 and D trees, while for *G+CRF19* data set it formed a sister-group to the G subtree (the *G+CRF19* data set contained few sequences and did not include G sequences branching before the split with the CRF19 clade). There were several mis-annotated sequences in CU data set. For example *CU1437-17* was placed within the D sequences in the *D+CRF19* tree, while annotated in CU as CRF19: As it only contained the D-part of the sequence (and therefore could not be distinguished between CRF19 and D during the non-phylogeny-based subtyping) the annotation was most probably erroneous. In a similar manner, the CRF19 clusters of *A1+CRF19* and *D+CRF19* trees included a few A1 and D sequences respectively: Those sequences only contained the A1 or D fragments and therefore were likely mistyped as well.

**Fig 1.**
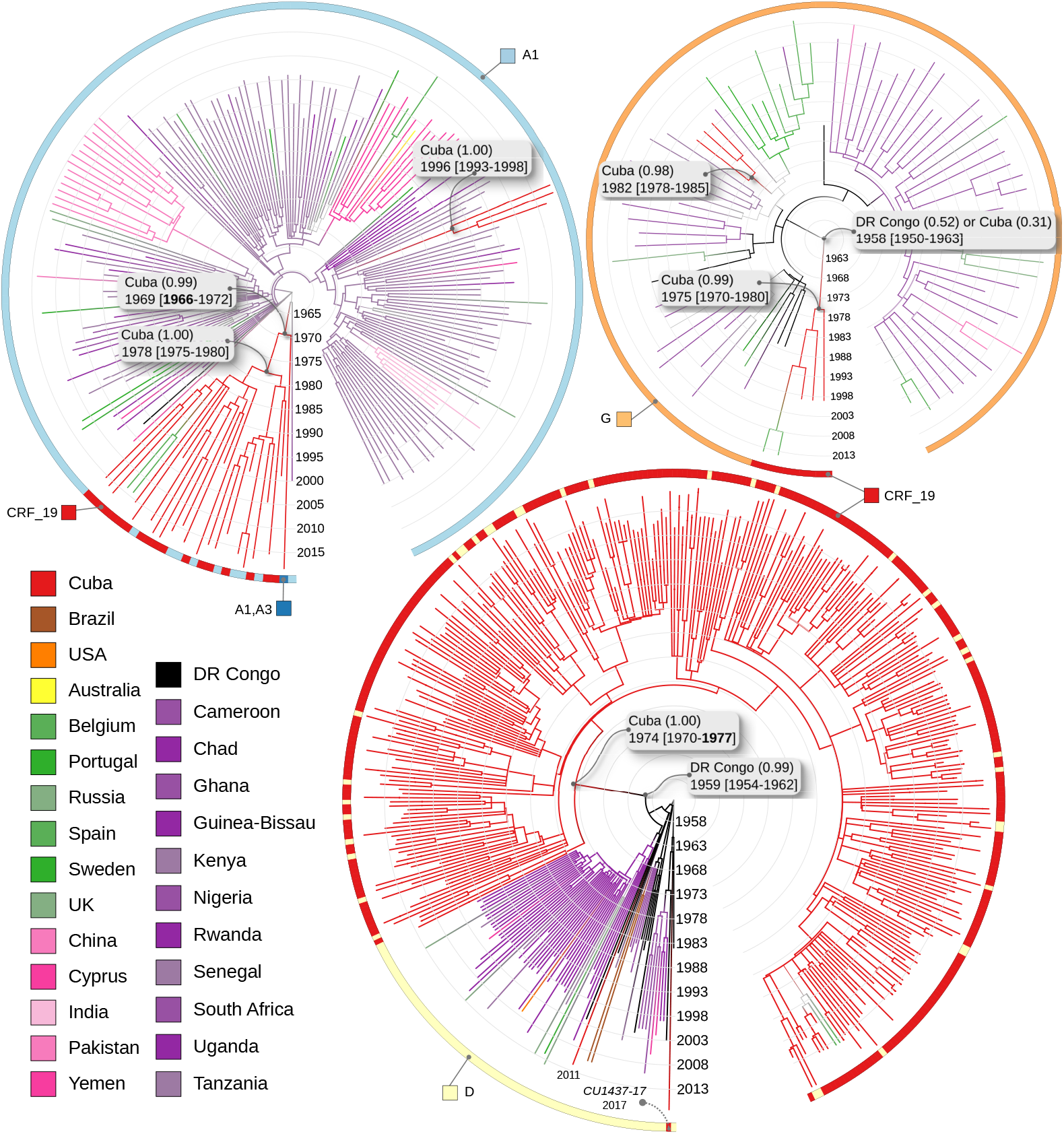
**Time-scaled trees** reconstructed for the three data sets: *A1+CRF19* (top left), *G+CRF19* (top right), and *D+CRF19* (bottom centre). Nodes and branches are coloured by country (ACR performed by PastML, MPPA+F81), as explained in the legend in bottom left: Cuba is coloured red, African countries are in shades of violet apart from the the DRC (black), European countries are in shades of green, Asian countries are in shades of pink, Australia is yellow. Colourstrips around the trees show sample subtypes: A1 (light-blue), D (yellow), G (light-orange), CRF19 (red), and A1-A3 recombinant (blue, note that this sequence was used in the *A1+CRF19* data set as its part that aligns with the A1-part of CRF19 is fully A1). CRF19 sequences form a cluster for all data sets. Subtype misannotations lead to several D- and A1-annotated sequences within the corresponding CRF19 clusters, and one CRF19-annotated sequence (*CU1437-17*) outside of the CRF19 cluster in the *D+CRF19* tree. Dates (with CIs, reconstructed by LSD2) and countries (with marginal probabilities, reconstructed by PastML) of the CRF19 cluster MRCAs and their non-CRF19 (i.e. A1, D or G) parent nodes are shown. The dates of non-recombinant Cuban clusters are also shown.

### Dating

The dates obtained for the MRCA of the CRF19 cluster and for its parent (i.e. the most recent D/A1/G-subtype ancestor of the CRF19 cluster) estimated on the three data sets are shown in Fig 1 and in Table 2. For all the data sets the CRF19 MRCA date is estimated in the 1970*s*, and the most recent A1/D/G-subtype ancestor dates are 1969*/*1959*/*1958 respectively. Note that as the three data sets include different CRF19 sequences, the CRF19 cluster MRCA dates do not have to agree, but they all correspond to the time after CRF19 recombination. Similarly, the A1/D/G-subtype ancestor dates correspond to the time before recombination and do not have to be the same for the three data sets, but have to be more ancient than all of the CRF19 MRCA dates. Based on our estimations, the dates of origin inferred by using different parts of the CRF19 are all consistent and go in the same direction: we obtain a confident estimation of the date of recombination, occurring between 1966 (lower CI of the date when in *A1+CRF19* data set it was still A1) and 1977 (upper CI of CRF19 MRCA in the *D+CRF19* data set). We also dated the non-recombinant A1/D/G-subtype Cuban sequences in our data sets. They were estimated to be introduced to Cuba later: 1982 for G, 1996 for A1 and in the 2010*s* for D (see Fig 1), supporting the hypothesis that the recombination occurred in Africa, as co-infection and thus high-level circulation of the 3 subtypes was needed for recombination.

**Table 2.** Dates and CIs of the MRCA of CRF19 and its non-CRF19 parent, etimated on different data sets.

| data set | CRF19 MRCA date | non-CRF19 MRCA date |
| --- | --- | --- |
| *D+CRF19* | 1974 [1972 – 1977] | 1959 [1954 – 1962] |
| *A1+CRF19* | 1978 [1975 – 1980] | 1969 [1966 – 1972] |
| *G+CRF19* | 1975 [1970 – 1980] | 1958 [1950 – 1963] |

### Phylogeography

The country predictions (and their marginal probabilities) for the MRCA of the CRF19 cluster and of its parent (i.e. the last A1/D/G-subtype ancestor) estimated on the three data sets are shown in Fig 1. The results suggest that CRF19 recombined in the Cuban community, as the CRF19 MRCAs in all data sets and the A1 ancestor have Cuba as the country reconstruction. The reconstructions also suggest that the D-strain ancestor has a DRC origin, while the G-strain reconstruction hesitates between Cuba and the DRC (with a slight preference for the DRC). Overall, these results fit well with the hypothesis of a recombination within the Cuban community in the DRC around 1970*s* [1966 – 1977].

We further performed a province-level analysis of the CRF19 subtree from the *D+CRF19* data set. The root of CRF19 subtree (dated in 1974) was unresolved with two most probable provinces being Villa Clara (0.64 as marginal probability) and La Habana (0.24). This prediction is not surprising and is consistent with respect to the previous dating and country predictions: the CRF19 subtree most likely originated in the Cuban community in sub-Saharan Africa. From there the epidemic spread to Villa Clara very early (marginal probability of 0.92, date 1977, CI: 1973 – 1979), before HIV discovery in the 1980*s* and likely before its introduction in a number of developed countries by the very end of 1970*s* (e.g. USA, UK [15]). Then CRF19 gave rise to multiple introductions from Villa Clara to La Habana in the 1980*s* and 1990*s* and then further spread to other provinces (Fig 2).

**Fig 2.**
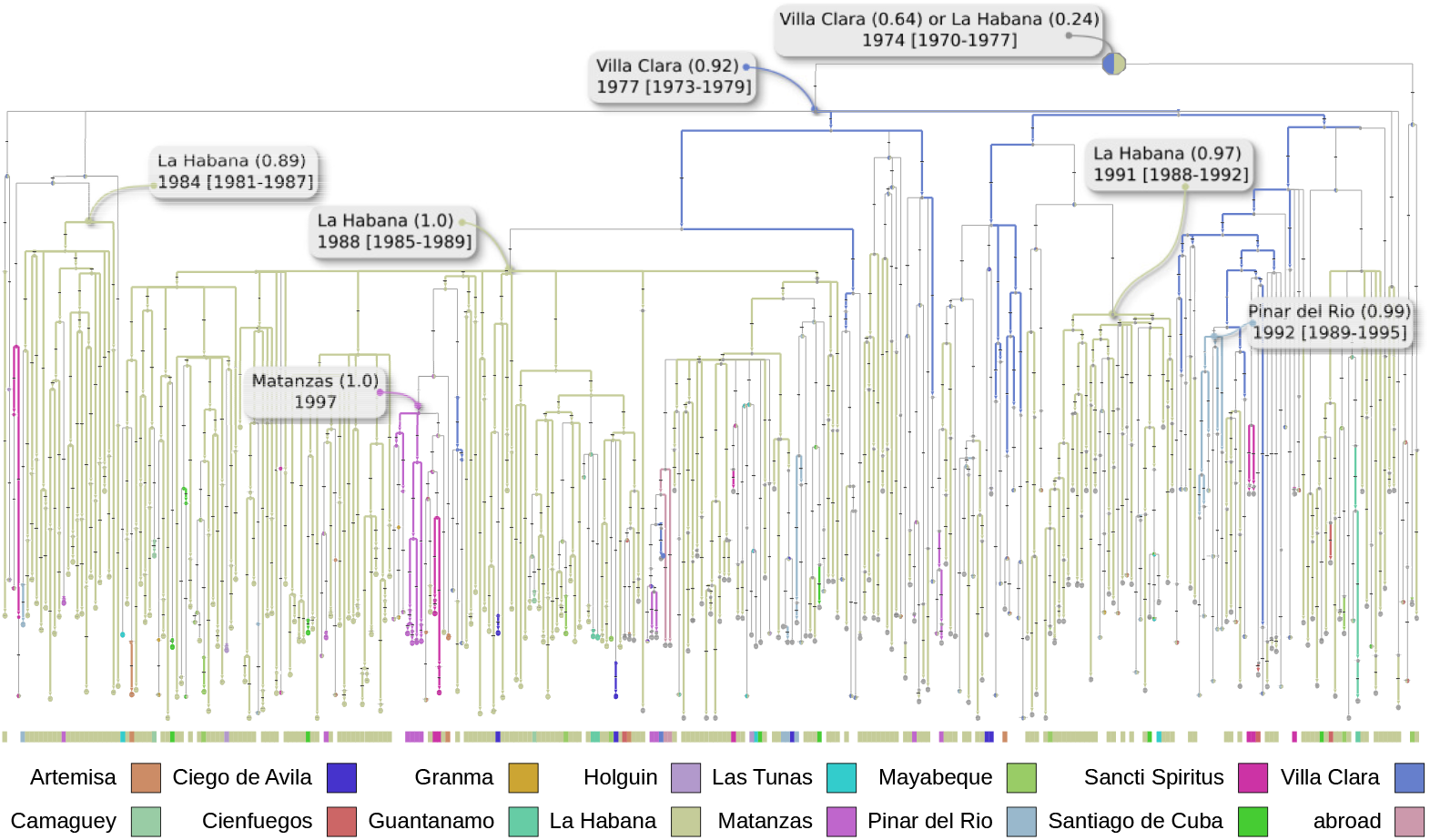
Phylogeography (province level) of CRF19. ACR and visualisation are done with PastML [16] (MPPA+F81). Nodes and branches are coloured by provinces (colour code is explained below), gray thinner branches correspond to a province change, multi-coloured nodes correspond to multiple possible states (e.g. the root, unresolved between Villa Clara and La Habana). The oldest resolved node and its date are indicated and correspond to Villa Clara (marginal probability 0.92), the dates and marginal probabilities of the main introductions from Villa Clara to La Habana are also marked.

### Transmission mode

ACR for the transmission mode (heterosexual (HET) or a transmission between MSM) reconstructed the most likely state of the origin of the epidemic to HET (marginal probability of 0.89 vs 0.11 for MSM), but suggested that in the 1980*s* it quickly spread to and within the MSM community and had multiple transmissions back to HET (often female): Out of 312 transmissions in our tree, 53 were within the HET community, 74 from HET to MSM, 155 within the MSM community, and 30 from MSM to HET (including 14 to females), see Fig 3 for more details.

**Fig 3.**
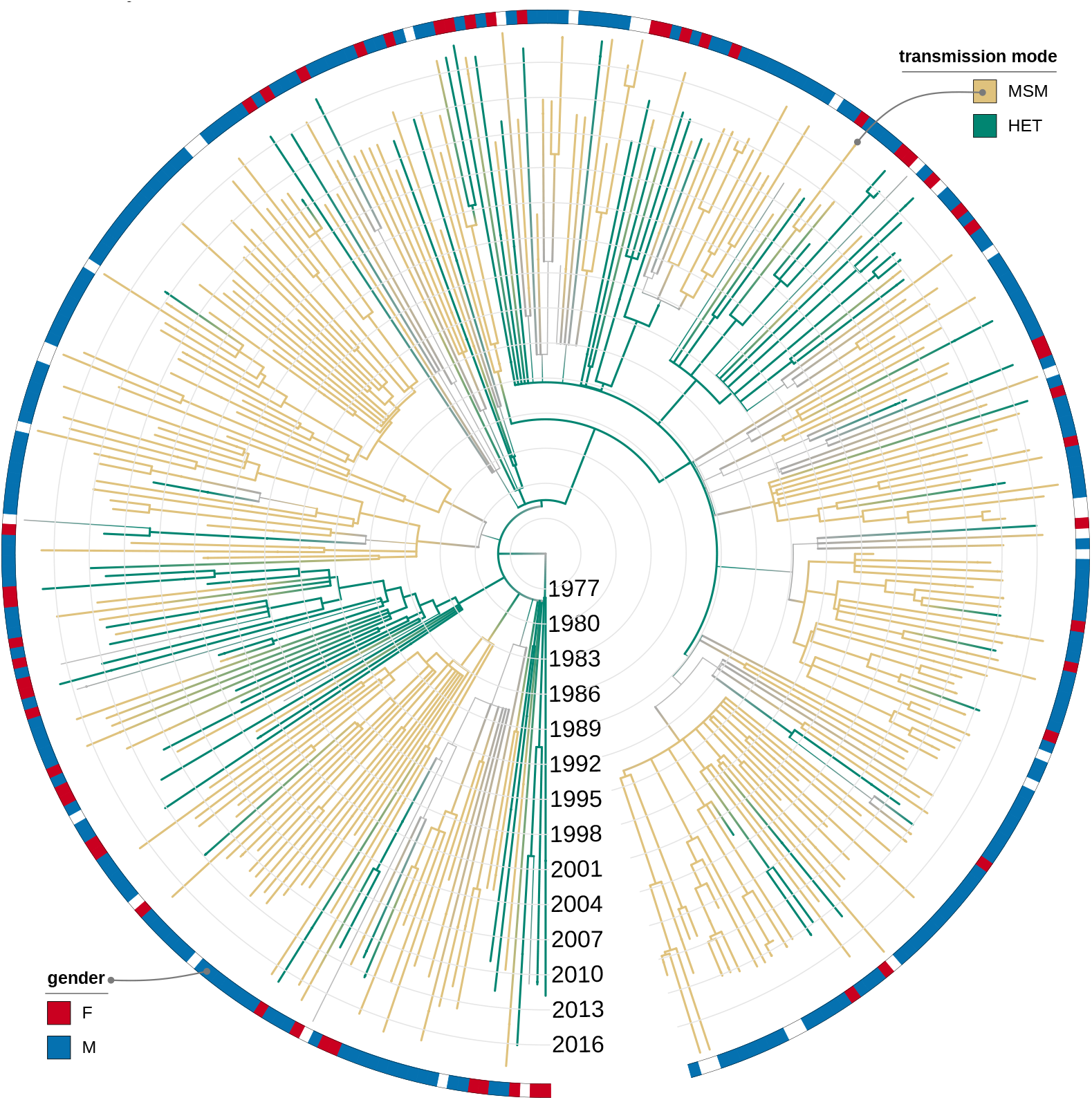
ACR for the transmission mode of CRF19. ACR is done with PastML [16] (MPPA+F81), the visualisation is performed with iTOL [17]. Nodes and branches are coloured by transmission mode (green for HET, beige for MSM), gray thinner branches correspond to a state change, gray nodes have unresolved states (MSM or HET). The colourstrip around the tree shows the gender of the sampled individuals (red for female, blue for male), and we see several transmissions from the MSM community to female HET individuals.

### Drug resistance

We analysed the presence/absence of DRMs with prevalence of at least 10%: RT:M41L, RT:T215Y, RT:K101E, RT:K219E, RT:K103N, RT:G190A, RT:D67N, RT:Y181C, and RT:M184V. For most of them hardly any TDR clusters were detected (the largest TDR cluster contained 2 patients), which is probably related to undersampling (CU data set contained about 4 9% of the estimated number of people living with HIV in Cuba by 2018, according to UNIAIDS: unaids.org/en/regionscountries/countries/cuba), and relatively recent introduction of the corresponding ARVs. The latter is supported by the fact that for RT:M41L some TDR was present (see Fig 4): RT:M41L is associated with resistance to AZT [18], the first ARV accepted in Cuba (used since 1987).

**Fig 4.**
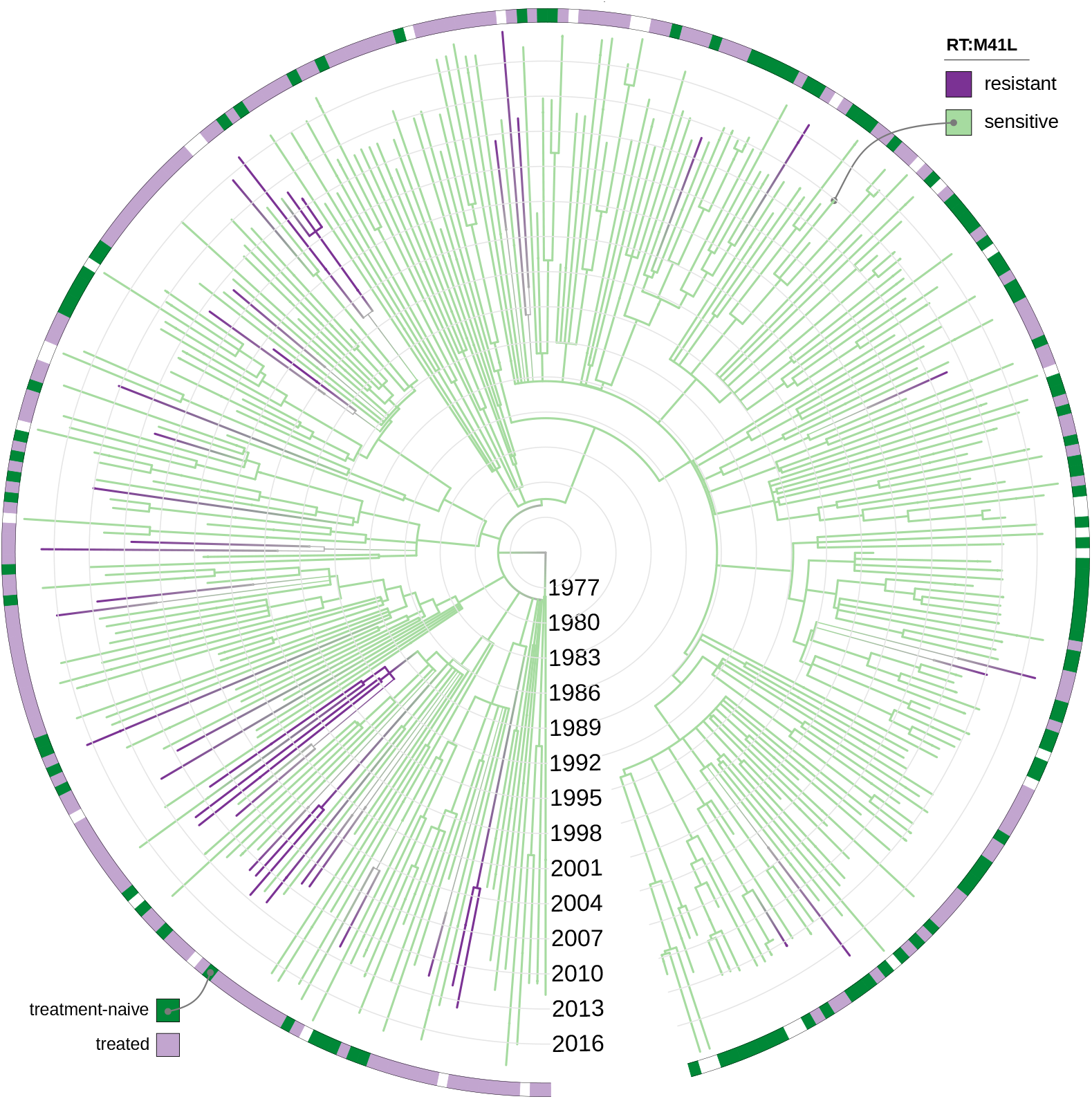
ACR for the presence of RT:M41L DRM in the CRF19-infected individuals. ACR is done with PastML [16] (MPPA+F81), the visualisation is performed with iTOL [17]. Nodes and branches are coloured by RT:M41L (violet for resistant (the mutation is present), light-green for sensitive). The colourstrip around the tree shows the treatment status of the sampled individuals (dark-green for treatment-naïve, lilac for treatment-experienced).

## Discussion

CRF19 is a predominant circulating recombinant form of HIV-1 in Cuba, with 24.1% of new HIV-positive patients being infected with it [19]. We investigated its origin and transmission patterns using new sequence data from Cuba.

Based on our date estimations, in 1966 CRF19 still did not exist as in the *A1+CRF19* data set it was still A1 (lower CI), but by 1977 it already recombined as in the *D+CRF19* data set this date is the upper CI of the CRF19 MRCA. This estimate is older than the one obtained by Delatorre and Bello [12] (CI: 1983 – 1991) but is not contradictory as Delatorre and Bello estimated the date of the CRF19 MRCA on a data set that contained less CRF19 sequences and hence should be regarded as the upper bound of the recombination date. Our result leaves both a recombination in Cuba and a recombination prior to introduction to Cuba possible (as Cubans were present in sub-Saharan Africa and especially DR Congo since the 1960s [5]). Recombination in Cuba was proposed by Delatorre and Bello for other prevalent HIV-1 recombinant forms (CRF20 BG, CRF23 BG, CRF24 BG), which probably emerged between 1996 – 1998 from a single CRF BG ancestor dated in 1991. However, for CRF19 it seems unlikely that the recombination event took place in Cuba considering that (1) we traced MRCA for CRF19 back to 1977 or before, (2) the first Cuban HIV-1 patients were not diagnosed until 1986, (3) the non-recombinant A1/D/G-subtype Cuban sequences in our data sets were estimated to be introduced to Cuba after 1977 (1982 for G, 1996 for A1 and in the 2010*s* for D), while (4) recombination had to occur in a scenario of co-circulation of subtypes A, D, and G. Moreover, countries from sub-Saharan Africa were proposed to be sites of origin for other AG and AD recombinants within the same period of time [20, 21]. All together, we can conclude that CRF19 was among the very first CRFs/subtypes to be introduced in Cuba, by the mid of 1970*s*. Other subtypes have undergone the same journey, from Central Africa to the Caribbean, for example subtype B, which was present in Haiti since the end of the 1960s [22]. However, while subtype B rapidly reached the USA and then Europe and the rest of the world, CRF19 stayed at Cuba, where it represents a major issue due to its prevalence and short time to AIDS.

To our knowledge, this is the first study which attempts to predict the origin of HIV-1 epidemic (and CRF19) by Cuban province employing sequence data. The phylogeography analysis detected that La Habana and Villa Clara played a major role in epidemic spread. We estimated that CRF19 was initially introduced to Villa Clara in the late 1970*s* (marginal probability of 0.92), had multiple introductions to La Habana in 1980*s* and 1990*s* and spread from there to other provinces. In agreement with our results, Dhanjal *et al.* [23] employed a series of graph algorithms to predict the evolution of the HIV/AIDS network in Cuba between the years 1986 and 2004. These authors suggested that at the end of 1986 epidemic was mostly located in La Habana, with 31 detections, and pointed out that Villa Clara and Sancti Spiritus were among the first provinces showing an increase in the detection rate. A sudden growth in detection in La Habana was expected given the high population density of this city and the difficulties to control the epidemic. In addition, an analysis performed by Miranda *et al.* [24] showed that, until 2010, Villa Clara remained among the provinces with a high incidence rate.

Our analysis on the transmission mode recreates some of the Cuban HIV-1 epidemic features such as a rapid spread among MSM community and multiple introductions to HET population (16% of all transmissions with MSM source, 50% of which were to females), indicating potential importance of bisexual individuals in the epidemic spread. According to epidemiological data, Cuban epidemic started as a mainly HET epidemic, after 1989 detection prevailed in homo/bi-sexuals, however, by 2002, a sudden rise in HET group occurred, ultimately showing a tendency to parity [24, 25]. In our reconstruction, CRF19 epidemic in Cuba followed a similar pattern and started in HET group, which agrees with the Pérez *et al.* [25] finding that among 28 individuals diagnosed with HIV between 1986 and 1990, 25 were HET and were infected with a high diversity of genetic forms.

Lastly, we did not detect almost any transmitted drug resistance, apart from several small TDR clusters for RT:M41L, a DRM which is associated with resistance to AZT, the first ARV accepted in Cuba (in 1987). One of the reasons why we could not trace almost any TDR might be because the sequences employed in this study were obtained as a result of Drug Resistance Surveillance in Cuba, and not the result of contact tracing studies. Several studies on the prevalence of TDR and a national survey from 2017 [19, 26] showed that, despite a high drug resistance incidence (30% pre-treatment drug resistance), no clusters of TDR have been identified in Cuba to date for any HIV-1 subtype. However, epidemiological data combined with a small genetic distance among sequences from different patients has evidenced that, indeed, transmission of resistant HIV-1 genetic variants occurs in Cuba. This was almost invisible in our data, most likely due to their scarcity (*<* 10% of infected individuals).

Considering all this, we suggest that CRF19 recombined in the Cuban community present in sub-Saharan Africa since the 1960s, most probably in the DRC, as a consequence of the co-circulation of subtypes A, D and G. It was introduced in Cuba very early, in the 1970s, and started to spread from a single transmission event, first in Villa Clara and then in the 1980s in La Habana. This early introduction to Cuba, which is earlier than the introduction of A, D and G subtypes, for example, could explain the local success of CRF19, which had no particular success in other parts of the world. This local success would thus result from a founder effect rather than some transmission advantage of CRF19, as supported by the relatively stable incidence of CRF19 through time: 18.3% in 2007 (sampled in 2003), 27.5% in 2012 (diagnosed in 2009 – 2010), 24.1% in 2019 (sampled in 2017). However, importantly, the fact that the time to AIDS with CRF19 is substantially lower than with other subtypes and CRFs (3 years versus 5 – 10 years [6]), makes it crucial to survey CRF19 sub-epidemics not only in Cuba but also in other parts of the world having regular exchanges with Cuba.

## Materials and methods

### Alignment and subtyping

LA dataset contained 3 884 aligned and subtyped sequences of all M-group HIV-1 subtypes and recombinants. CU contained 1 674 partial *pol* and 164 partial *env* sequences (of various subtypes) recently sampled in Cuba. We subtyped (pure subtypes and recombination positions) and aligned them against LA using jpHMM [27] (for detailed options see Analysis pipelines). When several sequences were present for the same patient in CU, we kept only the first one (in terms of sampling date). Most of the sequences in the CU data set were pre-annotated with a subtype; we replaced these annotations with jpHMM predictions in cases when they were contradictory (e.g. not matching jpHMM breakpoints for CRF19). All together, we obtained a large MSA of 5 707 sequences, including both LA and CU, from which we extracted our *D+CRF19*, *A1+CRF19* and *G+CRF19* data sets (see Table 1 for details).

### Metadata

LA alignment contained country and sampling date annotations. The sequences from CU were also annotated with the province and date of diagnostics (which sometimes preceded sampling by several years), patient gender, risk groups (HET or MSM) and treatment status (treated or treatment-naive). For the DRM analysis we extracted the presence/absence of the surveillance DRMs with Sierra web service [28] (for detailed options see Analysis pipelines)).

### Time-scaled trees

For each data set we reconstructed a phylogenetic tree using RAxML-NG (v0.9.0, evolutionary model GTR+G6+FO+IO, for detailed options see Analysis pipelines) [29] and rooted it with an outgroup (5 LA sequences of a different subtype, B for *D+CRF19*, A6 for *A1+CRF19*, A for *G+CRF19*, which were then removed).

We then dated each tree with LSD2 [30] (v1.4.2: github.com/tothuhien/lsd2/releases/tag/v.1.4.2, under strict molecular clock with outlier removal, for detailed options see Analysis pipelines) using tip sampling dates, and constraints based on the diagnostics dates. As a phylogenetic tree is a proxy for a transmission tree with incomplete sampling, its branches correspond to patients and its internal nodes correspond to virus transmissions from a donor patient (parent branch and one of the child branches) to a recipient patient (the other child branch). As the recipient patient must have been infected (internal node) before being diagnosed, we put constraints for the dates of the internal nodes that had two tip children, at the dates of diagnostics of their recipients. As one cannot distinguish between the recipient and the donor branches in the tree (see Fig 5 for more details), we picked the more recent diagnostics date between the two and imposed the transmission node date to be before this diagnostics date. The temporal constraints on internal nodes are incorporated in LSD2 using the active-set method in the same way as it was used in LSD [30] to ensure the temporal precedence on ancestral nodes.

**Fig 5.**
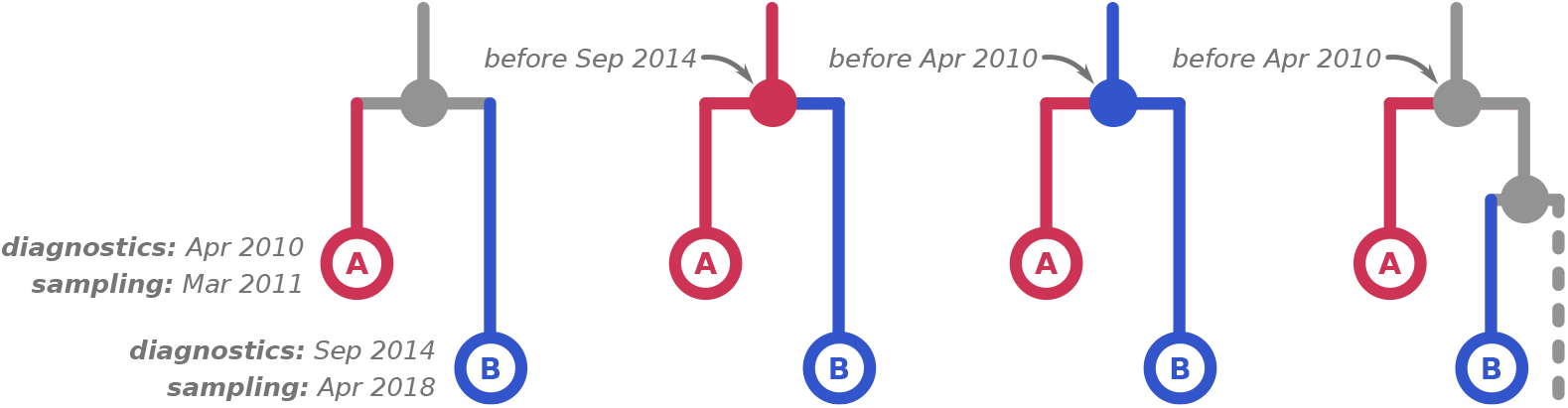
Possible transmission scenarios for two samples. An example internal node that has two tip children, *A* and *B*, and their sampling and diagnostics dates are shown on the left. As the internal node corresponds to a transmission, three scenarios are possible: (middle left) *A* is the donor, therefore the transmission must have happened before the recipient (*B*) got diagnosed; (middle right) *B* is the donor, therefore the transmission must have happened before the recipient (*A*) got diagnosed; (right) the donor was another individual that was not sampled and does not appear in the tree, therefore that transmission must have happened before both *A* and *B* got diagnosed. In practice, however, we cannot know which of the three scenarios is the correct one and therefore have to pick the more recent diagnostics date (*B*, Sep 2014) as the upper-bound constraint for the internal node.

### Ancestral character reconstruction (ACR)

We analysed geographic origin and transmission patterns of the CRF19 epidemic worldwide and in Cuba using ACR with PastML tool (v1.20, for detailed options see Analysis pipelines) [16]. PastML infers an ancestral scenario for a given character (e.g. location) on a rooted phylogenetic tree, by assigning the likely ancestral character states to every node in the tree, based on the character annotations provided for some of the tree nodes (typically tips). PastML implements a very fast ACR in maximum likelihood framework, under an *s*-state (where *s* is the number of possible character states, e.g. countries) F81-like [31] model, and can handle very large phylogenies (dozens of thousands of tips), while being robust to mild sampling bias and model violations [16]. With marginal posterior probabilities approximation (MPPA) method a unique state is predicted in tree nodes with low uncertainty, whereas several states are predicted if they have similar marginal probabilities.

For phylogeographic analysis we reconstructed ancestral scenarios for country annotation on the full trees reconstructed from the three data sets. Additionally, we reconstructed ancestral characters for Cuban provinces on the CRF19 subtree of the *D+CRF19* tree (which included most of the CRF19 sequences in our data sets). As for the province-level analysis the annotations were specified at the date of diagnostics, which often preceded sampling, we created additional internal nodes in the tree, corresponding to diagnostics, named them, and associated the input province annotations for PastML with them instead of tree tips (see Fig 6). To add the diagnostics nodes, we processed each tip *T* whose patient’s diagnostics date *diag*(*T*) preceded the sampling date *d*(*T*) *> diag*(*T*). If the diagnostics happened after the last transmission (i.e. its parent node *P*, where *d*(*P*) *< diag*(*T*)), we split the tip branch into two parts with a one-child internal node *D* at the time of diagnostics (*d*(*D*) = *diag*(*T*)): The branch length of *D* became *dist*(*D*) = *diag*(*T*) − *d*(*P*), while the branch length of *T* became *dist*(*T*) = *d*(*T*) *diag*(*T*). Otherwise (when the diagnostics happened before the last transmission *diag*(*T*) *< d*(*P*), which meant that the parent branch corresponded to the same (donor) patient as the tip), we processed the parent node (or recursively the grandparent node, etc.) instead, in a similar manner.

**Fig 6.**
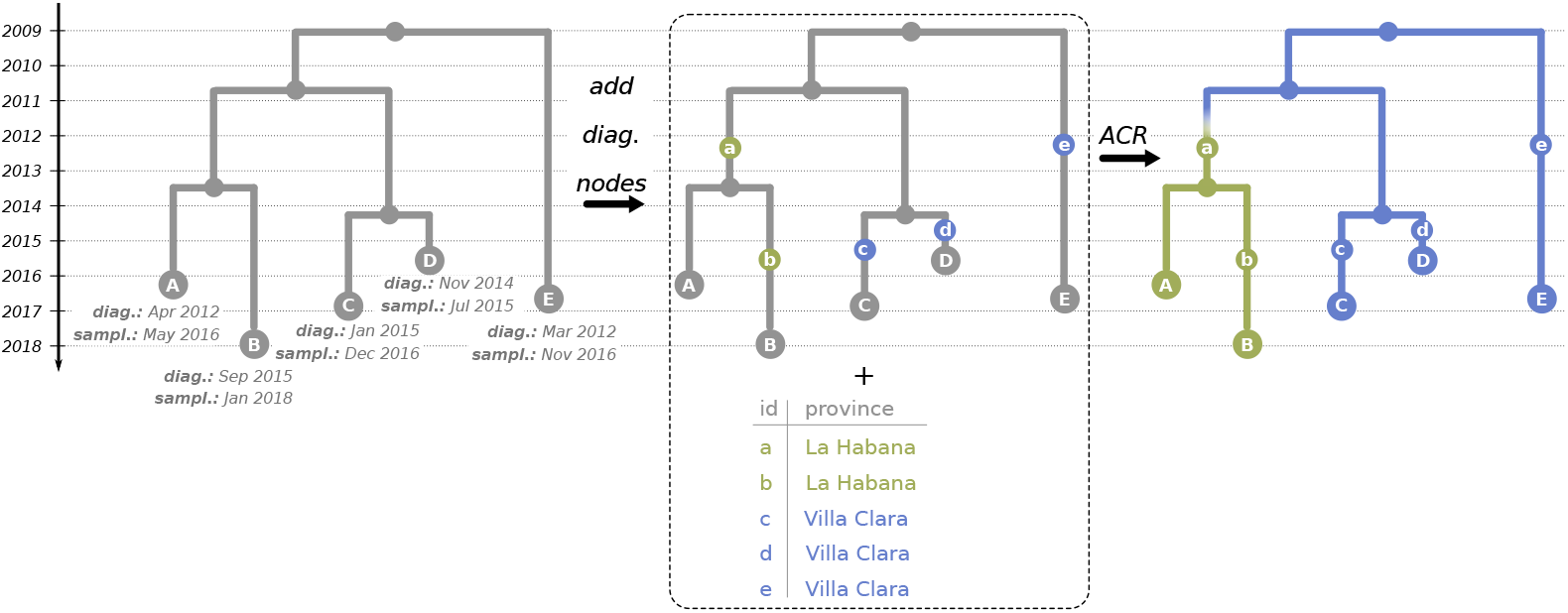
ACR for provinces of Cuba. To reconstruct the ancestral states for characters annotated at diagnostics dates (such as provinces of Cuba), we added additional nodes to the time-scaled trees at times of diagnostics, named those nodes (on the middle panel, the diagnostics nodes, e.g. *a*, correspond to the tips with the same names but capitalised, e.g. *A*), and associated the input annotations for PastML with them. We then reconstructed ancestral and tip characters with PastML based on those annotations (see text for calculation details).

PastML allows for input annotations to be associated with any node in the tree (not only tips), and infers the states of the other nodes (including unannotated tips) during ACR. The computation of the Up and Down-likelihoods for a one-child node *D* being in a state *i* is a corner case of the formulas described in the “Materials and Methods” section of the original PastML publication [16]: *Down*(*D, i*) = Σ_*j*_ *PC*(*i → j/dist*(*T*))*Down*(*T, j*), *U*_*p*_(*D, i*) = Σ_*j*_ *PC*(*i → j/dist*(*D*))*U*_*p*_(*P, j*). If an internal node’s state is provided in the input annotation file, then the Down- and Up-likelihoods of it being in any other state are reset to 0. The rest of the formulas from [16] are directly applicable.

To further understand the Cuban CRF19 epidemic, we also reconstructed ancestral scenarios for transmission mode (i.e. MSM or HET). The patients in the CU data set were annotated with their risk group (MSM or HET), and we used these data as a proxy for their potential HIV transmission mode at the moment of diagnostic and sampling: We associated the same annotations to both the corresponding single-child internal nodes (representing diagnostics) and to the tips (representing sampling). Consistently, the transmission mode is the same for the donor and the recipient at the moment of transmission (internal nodes of the tree), but can change with time for the same person. Note, moreover, that the MSM/HET status is uncertain, as some MSM might be bisexual or declared as HET [32].

Lastly, we analysed DRM presence/absence on the CRF19 subtree of the *D+CRF19* tree as the D-part of CRF19 includes the parts of the virus genome (PR, RT) that may contain DRMs. We analysed the DRMs that had a prevalence *>* 10%: RT:M41L, RT:D67N, RT:K101E, RT:K103N, RT:Y181C, RT:M184V, RT:G190A, RT:T215Y, and RT:K219E. To reconstruct the ancestral character states for each DRM, we used a procedure visualised in Fig 7. We checked the antiretroviral drugs (ARVs) that can provoke the DRM and the dates of their introduction to Cuba (e.g. RT:M41L is associated with resistance to AZT, which was the first ARV for HIV and was used in Cuba since 1987, to D4T and potentially to DDI, both of which were used in Cuba since 2001). We then cut the time-scaled tree (with named internal nodes) at the earliest of these dates (e.g. 1987), therefore obtaining the pre-treatment-introduction tree and a forest of post-treatment-introduction subtrees. For the trees in the forest we added additional one-child root nodes (as parents of the corresponding tree roots, at distances that corresponded to the differences between the root dates and the ARV introduction date), which we marked as sensitive in the PastML input annotation file. We performed the ACR with PastML on the forest (functionality that we added to PastML since version 1.9.19): the forest likelihood is calculated by multiplying the likelihoods of all its trees. For the new roots their Up-likelihood was set to 1 for the sensitive state and to 0 for the resistant state, and the Down-likelihood calculated as described above. Once the forest ACR was performed we combined it with the all-sensitive annotations for the pre-treatment-introduction tree nodes.

**Fig 7.**
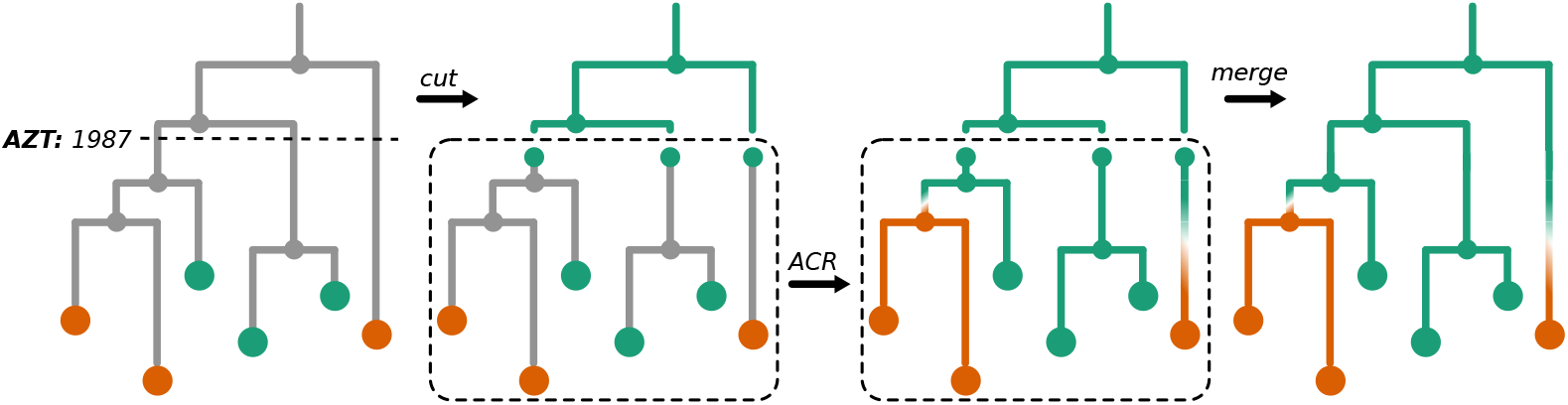
ACR for DRMs. To reconstruct the ancestral character states, resistant (violet) or sensitive (light-green), for a DRM (e.g. RT:M41L), we cut the time-scaled tree at the date of introduction to Cuba of the first ARV (as, for example, AZT for the DRM RT:M41L, used in Cuba since 1987) that can provoke this DRM (left panel), obtaining the pre-treatment-introduction tree (upper part of the tree) and a forest of post-treatment-introduction subtrees (bottom part). For the trees in the forest we then marked their roots as sensitive (middle left panel). We performed the ACR with PastML on the forest (middle right panel) and combined the results with the the all-sensitive annotation for the pre-treatment-introduction tree nodes (right panel).

### Analysis pipelines

Snakemake [33] pipelines and ad hoc Python3 scripts used for the analyses described above are available on github.com/evolbioinfo/HIV1-CRF19_Cuba. Along with the subtyping, tree reconstruction, dating and ACR tools mentioned above, we used goalign (v0.3.1, github.com/evolbioinfo/goalign) for basic sequence alignment manipulations (counting the number of sequences and the alignment length, removing positions containing only gaps, etc.), as well as gotree (v0.3.0b, github.com/evolbioinfo/gotree) and ETE3 [34] for basic tree manipulations (format conversion, pruning, etc.).

## Acknowledgments

The authors would like to thank Dr. Gonzalo Bello for his helpful advice on phylogeography of CRF19. This work was supported by the Campus France collaborative programme “Partenariat Hubert Curien franco-cubain Carlos J. Finlay”. JV was supported by Ecole Normale Supérieure Paris-Saclay and by ED Frontières de l’Innovation en Recherche et Education, Programme Bettencourt.

